# The social dynamics of collective problem-solving

**DOI:** 10.1101/771014

**Authors:** Kyanoush Seyed Yahosseini, Mehdi Moussaïd

## Abstract

When searching for solutions to a problem, people often rely on the observation of their peers. How does this process of social learning impact the individual and the group’s performance? On the one hand, research has shown that individuals benefit from social learning in numerous situations and across many domains. Through social learning, individuals can access good solutions found by others, improve them, and share them in turn. On the other hand, this individual benefit may come at a cost: An excessive tendency to copy others often decreases the overall exploration volume of the group, thus reducing the diversity of discovered solutions, and eventually impairing the collective performance.

Here we investigate the conditions under which social learning can be beneficial or detrimental to individuals and to the group. For that, we model problem-solving as a search task and simulate various amounts of social learning. We avoid model specific considerations by relying on a simple framework whereby individuals gradually explore the search environment – a two-dimensional landscape of solutions – while being attracted to the best solution of the group.

Our results highlight a *collective search dilemma*: When group members learn from one another, they tend to improve their own individual performance at the expense of the collective performance. How is this dilemma affected by the structure of the search environment? By varying two structural aspects of the search environment, our results reveal that the negative effect of the dilemma is mitigated in more difficult environments.

Finally, we show that single individuals can profit from a high propensity of social learning, which in turn is damaging for the other group members. As a consequence, if individuals continually adapt their behavior to maximize their own payoff, groups converge to a sub-optimal level of social learning. Unraveling these intricate social dynamics helps to understand the complex picture of collective problem-solving.

## 1 Introduction

Humans and other social animals seldom solve problems in isolation. When searching for a solution, individuals often rely on the observation of their peers to decide where to explore, and when to exploit a previously discovered solution (Laland, 2004; Danchin et al., 2004; Goldstone et al., 2013). Social learning, or the act of learning by observing the solutions of others, has been shown to be beneficial to individuals in numerous situations and across many species (Rendell et al., 2010; McGrath, 1984; Galef and Laland, 2005; Boyd et al., 2011). Social learning enables good solutions to spread between individuals (Mason, Jones, et al., 2008), who can further improve them and share their refined solutions with others (Mason and Watts, 2012). Recent studies have demonstrated that collective search can succeed in solving highly complex optimization problems, such as reconstituting protein structures (Cooper et al., 2010), mapping neurons connectivity (Kim et al., 2014), or improving quantum transport (Sørensen et al., 2016).

However, research has also shown that under certain conditions, relying on social learning can have negative consequences at the collective level (Giraldeau et al., 2002; Laland, 2004). An excessive tendency to copy others can decrease the overall exploration volume, reduce the diversity of discovered solutions and thus impair the group’s performance (Lazer and Friedman, 2007; Lorenz et al., 2011). In particular, in “rugged” search environments characterized by numerous local optima, the risk that the group converges to a suboptimal solution is increased (Barkoczi and Galesic, 2016) – a phenomenon labeled maladaptive herding (Toyokawa et al., 2019), pre-mature convergence (Pandey et al., 2014), or negative information cascades (Bikhchandani et al., 1998).

These two opposed effects of social learning have sparked rich interdisciplinary research between cognitive sciences, biology, and economics (Lamberson, 2010; Bernstein et al., 2018; Hills et al., 2015; Berdahl et al., 2013). Most research agrees that a high collective performance requires a delicate balance between social learning and independent individual search (Bernstein et al., 2018; Rendell et al., 2010; Toyokawa et al., 2019; Derex, Perreault, et al., 2018). Yet, it remains unclear what social dynamics drives this phenomenon, how the structure of the search environment influences it, and what would be the optimal strategy to adopt – for the group and for the individuals.

For instance, one inconsistent aspect in extant research concerns the measurement of performance: whereas some studies rely on the average payoff of the group members (Mason, Jones, et al., 2008; Lazer and Friedman, 2007; Barkoczi and Galesic, 2016), others focus on the best solution found by the group (Cooper et al., 2010; Kempe and Mesoudi, 2014; Derex, Perreault, et al., 2018). In fact, these two measurements assess different aspects of the group dynamics: the average performance indicates how much group members can *individually* benefit from social learning, while the best solution is an indicator of the group’s *collective* performance. Here, we rely on numerical simulations to address two specific research questions: (1) How does the amount of social learning impact the individual and the collective performances?, and (2) how do the search environment and the social environment influence what is best for the individual and what is best for the group?

We model problem-solving as a search task in which individuals explore a vast, two-dimensional landscape of solutions, while assuming that each group member is trying to maximize its personal payoff (Barkoczi and Galesic, 2016; Lazer and Friedman, 2007). In order to avoid model-specific considerations, we use a generic model where individuals rely on a simple hill-climbing exploration strategy, while being attracted by the best solution of the group (Yahosseini and Moussaïd, 2019; Barkoczi and Galesic, 2016). We first study how the amount of social learning impacts the individual and the collective performance of the group, and highlight the existence of a strong tension between the two measurements. Second, we study the influence of the search environment, that is, the structure of the solution space, on both performance measurements. Third, we take a prescriptive view-point and investigate how much an individual should rely on social learning to maximize its own payoff, considering the behavior of its peers. Finally, we propose an approach to explain how a certain level of social learning could have developed through repeated social interactions.

## 2 Results

### 2.1 The collective search dilemma

The performance of a group can be assessed in two ways: (1) By evaluating the average payoff of each group member, the *individual performance*, and (2) by evaluating the payoff of the best solution that has been discovered by the whole group, the *collective performance*. We first investigate how individual and collective performances depend on the amount of social learning. For that, we introduce a social learning parameter *S* describing the degree to which an individual’s exploration is influenced by social information. Formally, *S* expresses the fraction of a simulation run where the individual is influenced by the solution of its peers rather than searching independently (see the model description in the method section). For instance, a social learning level of *S* = 0.8 indicates that the individual searches independently during the first 20% of the simulation run and relies on social learning for the remaining 80%.

As shown in figure 1A, an increase of social learning causes opposed effects on the two performance measurements: It increases the individual performance (up to a maximum for *S*_*opt*_ = 0.733), while at the same time decreases the collective performance. This conflicting effect reflects the collective search dilemma (Bernstein et al., 2018): When group members are influenced by each other, they improve their own individual performance at the expense of the collective performance.

**Figure 1:**
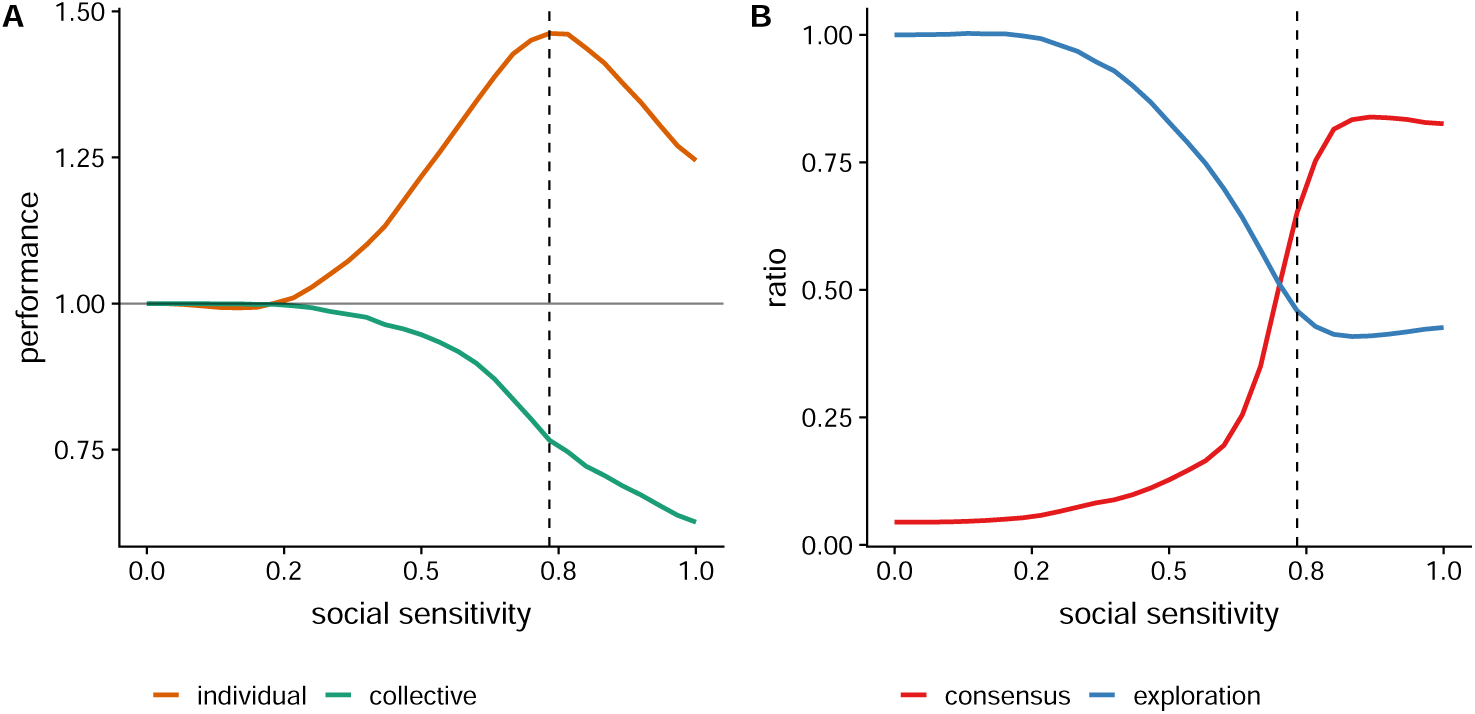
The collective search dilemma. (A) Impact of social learning on the individual (orange) and the collective (green) performance, relative to a group of independent searchers. Social learning yields an increase in individual performance at the expense of the collective performance. Individual performance is maximized for a learning level *S*_*opt*_ = 0.73 (dashed line). (B) The dilemma results from a drop in exploration (in blue; measured as the ratio of explored solutions relative to a group of independent searchers) and a boost of group consensus (in red; measured as the ratio of solutions shared by all agents). The best individual performance is achieved when exploration and group consensus are well-balanced.

We use the simulation results to better understand the dynamics of this dilemma. First, why is social learning improving individual performances? Having access to social information allows all individuals to profit from the best solution discovered by any of the other group members (figure 1B) (Bernstein et al., 2018; Mesoudi, 2011). Second, why is social learning hindering collective performance? An increase in social learning reduces the total number of explored solutions (figure 1B). As individuals are attracted by each other, they inevitably tend to explore the same region of the environment. Hence the group explores less and is thus less likely to find a good solution than when searching independently.

Figure 1A additionally highlights an optimal social learning level *S*_*opt*_ = 0.733 at which the individual performance is maximized. Excessive social learning (for *S > S*_*opt*_) causes a *premature convergence* effect whereby the agents explore too little and converge too early on the first discovered solutions – missing out potentially better ones. This effect lowers both the individual and collective performances (the individual and collective performances of a group of fully social agents with *S* = 1 are reduced by 11.7% and 16.5% compared to a group with social learning *S*_*opt*_).

## 3 Impact of the search environment

For now, we have assumed that the size and distribution of the peaks in the landscape (i.e., the local optima) are unstructured. In such a search environment, all peaks are evenly distributed and their payoffs are uncorrelated with their width. We introduce two continuous dimensions to study the effects of structural changes of the search environments: the kindness *k* and the dispersion *d* (see figure 2A). Kindness *k* manipulates the correlation between a peak’s width and payoff. For a negative correlation, higher peaks tend to have a smaller width (the environment is low in kindness) whereas for a positive correlation higher peaks tend to have a wider width (the environment is high in kindness). Dispersion *d* determines the structure of the locations of the peaks. For low *d*, peaks are more concentrated in one area (the environment is low in dispersion), while for high *d* they are more evenly distributed over the search environment (the environment is high in dispersion).

**Figure 2:**
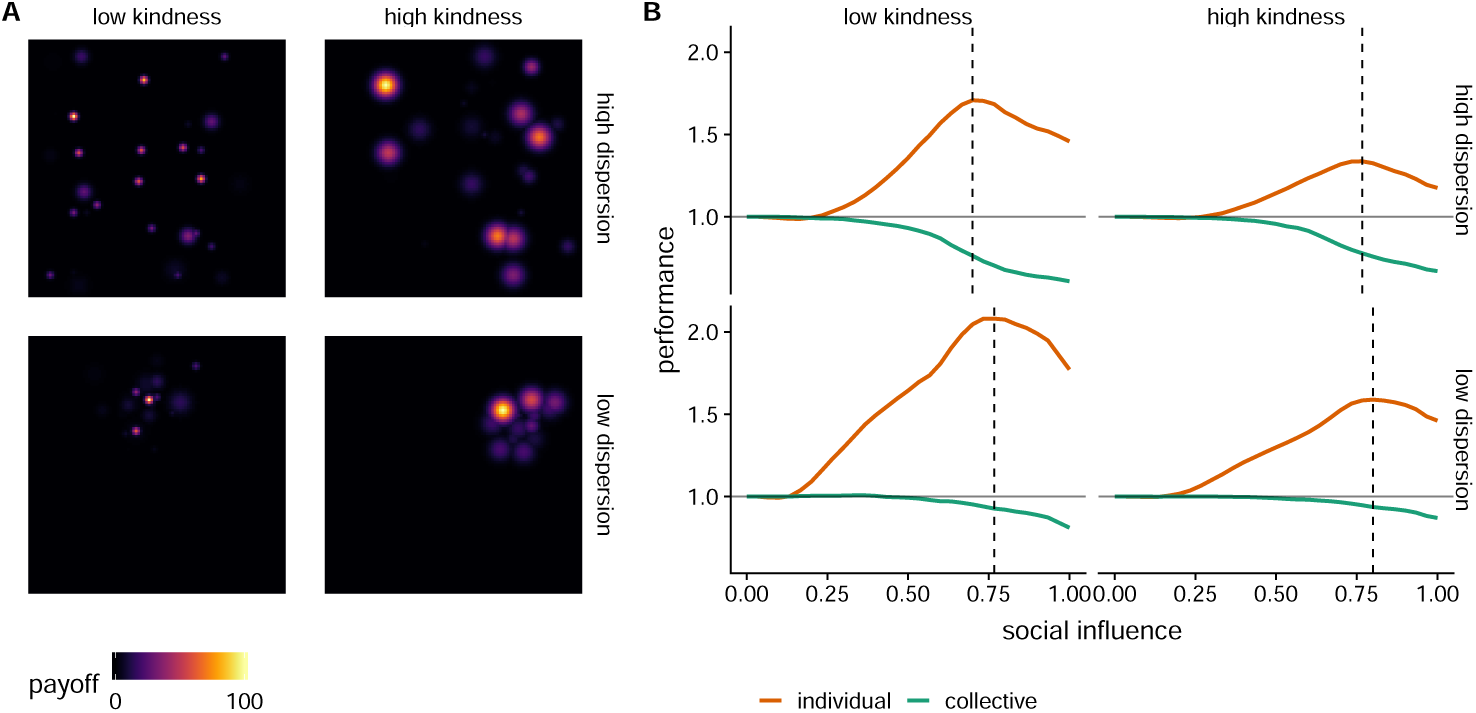
The structure of the search environment. (A) Illustrative examples of environments with low and high levels of kindness *k* (as columns), and low and high levels of dispersion *d* (as rows). Kindness refers to the correlation between the height and the width of the peaks. Dispersion determines the extent to which the peaks are clustered in the same area or scattered across the environment. (B) Corresponding patterns of individual and collective performances for different search environments. The search environments are of the same type as those illustrated in (A). Performances are relative to groups of independent searchers. The dashed vertical lines indicate the social learning levels *S*_*opt*_ that maximize the individual performance for each type of environment.

How does the structure of the search environment impacts the performances of a group? We find that the social search dilemma and the premature convergence persist across different physical environments, but change in amplitude (figure 2B). Overall, the beneficial effect of social learning on individual performances is larger in less kind environments, because social learning facilitates the discovery of hidden good peaks by the entire group (figure 3A). In addition, the negative effect of social learning on collective performance is smaller in when dispersion is lower (figure 3B). If no agent has found a satisfying peak at all, which is common when dispersion is low, social learning does not impair exploration and, thus, the collective performance.

**Figure 3:**
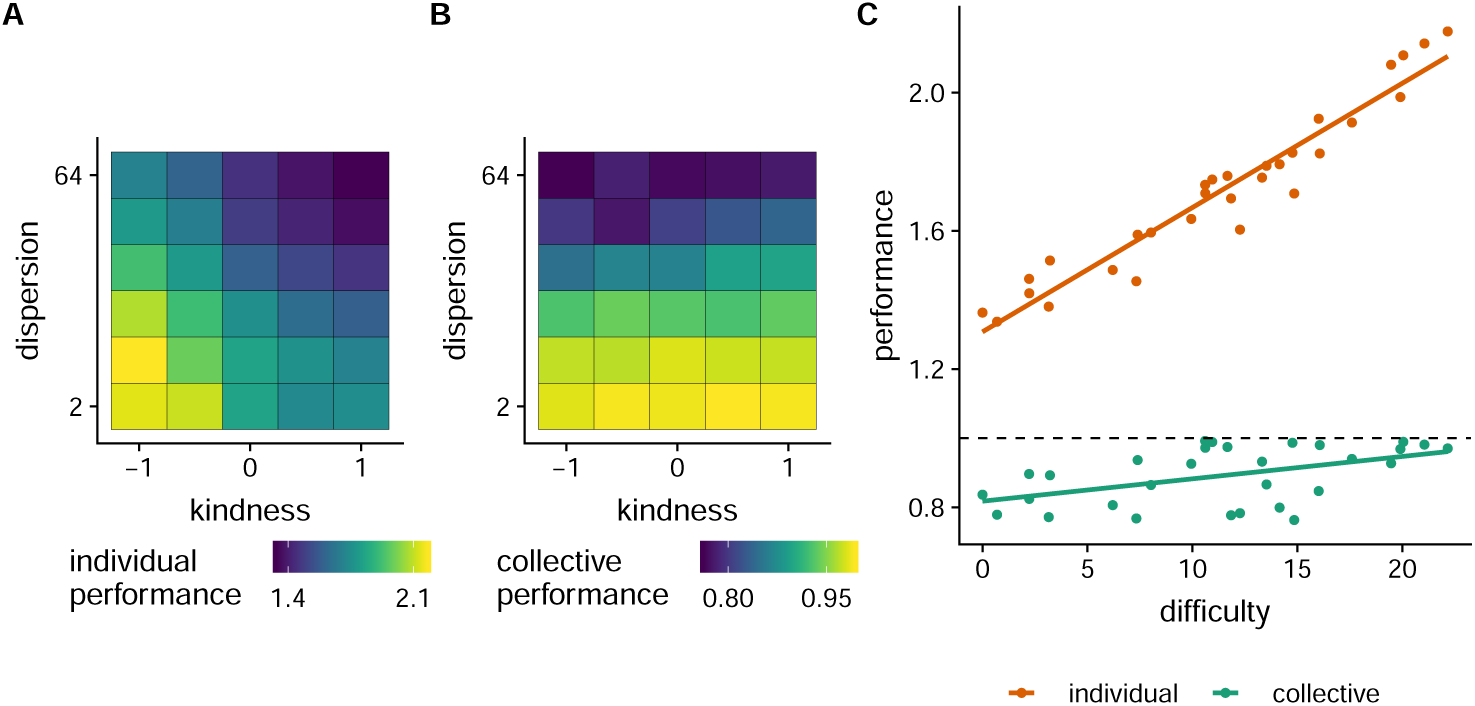
(A) Individual performances and (B) collective performances as the search environment varies in kindness (from low kindness *k* = − 1 to high kindness *k* = 1) and dispersion (low dispersion *d* = 2^1^ to high dispersion *d* = 2^6^). See the method section for more details. The color-coding indicates the performances at the optimal social learning levels *S*_*opt*_ (see dashed lines in figure 2B). (C) Individual and collective performances as a function of the difficulty level. Difficulty is measured as the inverse average individual performance of a group of independent searchers (higher is more difficult). Each dot represents the performances for a given combination of kindness and dispersion, taken at the optimal social learning level *S*_*opt*_. The dashed line indicates the performance of a group of independent searchers in that environment. Individual and collective performances increase with difficulty (linear model *r*^2^ = 0.93 and 0.25 for individual and collective performances, respectively).

Further manipulations of kindness and dispersion reveal a correlation between the difficulty of the environment, i.e. how well individual searchers solve it, and the beneficial influence of social sensitivity (see figure 3C). We find that, for more difficult environments, social learning leads to a bigger increase in individual performance and a smaller decrease in collective performance. Consequently in the most difficult search environments, the social search dilemma is mitigated.

### 3.1 Impact of the social environment

The environment is not limited to the search environment, but also concerns the social environment. How much should an individual rely on its peers, given the peer’s own social learning level? To answer this question, we systematically varied the social learning level for one individual in the group and for the rest of the group independently. That is, all agents except one are homogenous and share the same social learning level.

We find that the payoff of an individual highly depends on its social environment (figure 4A). From the perspective of an individual, the best performance is reached when all peers are independent searchers. In this case, the agent will be able to learn from others without undergoing the risk of premature convergence. On the contrary, social environments where other people have a high social sensitivity are the worst because the peers copying each other will reveal few solutions.

**Figure 4:**
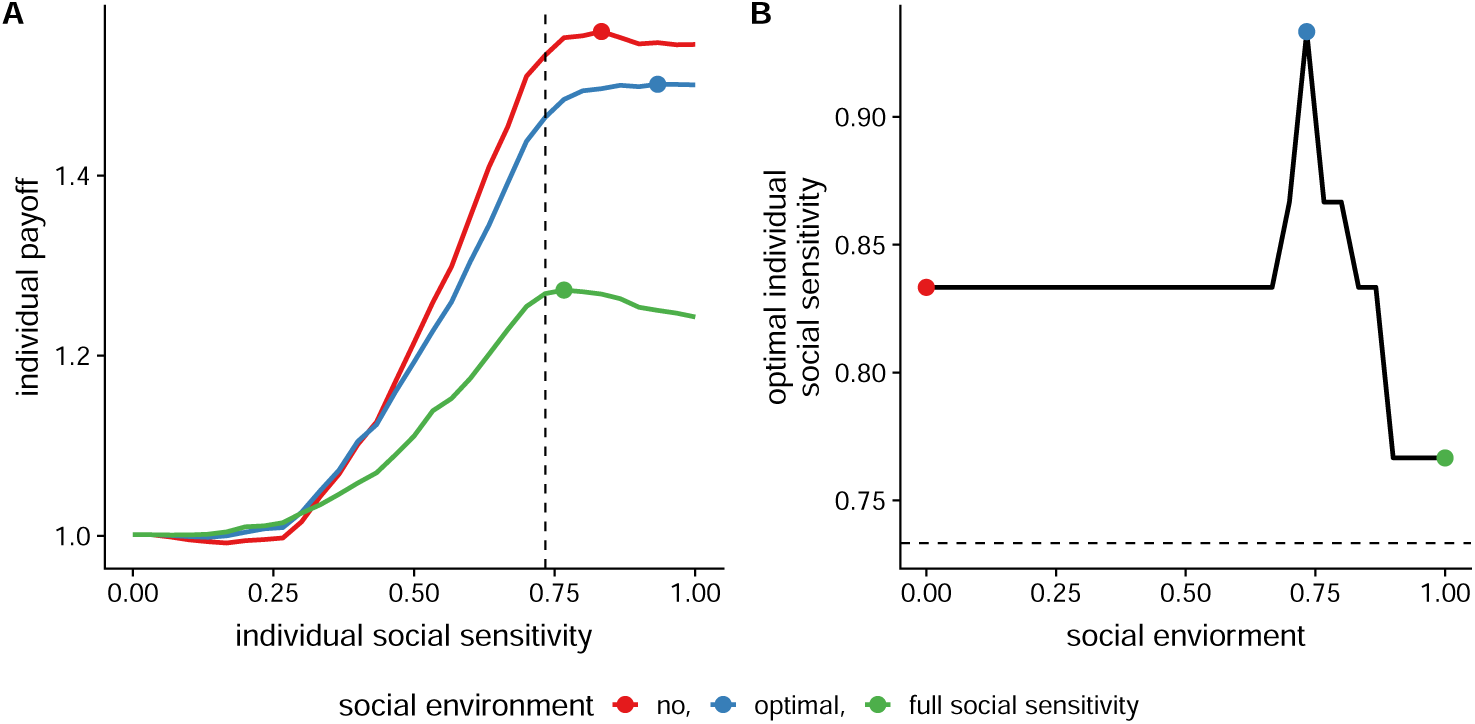
Influence of the social environment, measured as the level of social learning of one’s peers. (A) Impact of one agent’s social learning level on its payoff in different social environments (all other agents share the same social sensitivity, color coded). Dashed line indicates the optimal social learning for the group *S*_*opt*_. Points mark the curves maxima. (B) The social learning level maximizing an individual’s payoff in different social environments. These learning levels are consistently above the optimal group level *S*_*opt*_ (dashed line). Colored points indicate the social environments shown in A.

Our results also reveal that, in order to maximize its own payoff, an individual should rely on more social learning than the optimal level *S*_*opt*_ highlighted in figure 3B. Hence, what is optimal for an individual is different from what is optimal for the group, which might further increase the risk of premature convergence.

To study this hypothesis, we set up a simple model in which agents gradually adapt their individual social learning level *S* to maximize their own payoff (see figure 5 and method section). As expected, the average level of social learning increases slightly above *S*_*opt*_. That is, each individual is better off copying others more than what is optimal for the group, which leads to a decrease of performance when everybody does so. Hence, this model can explain why individuals often over rely on social information and account for the behavioral origins of premature convergence in some natural systems (Pandey et al., 2014).

**Figure 5:**
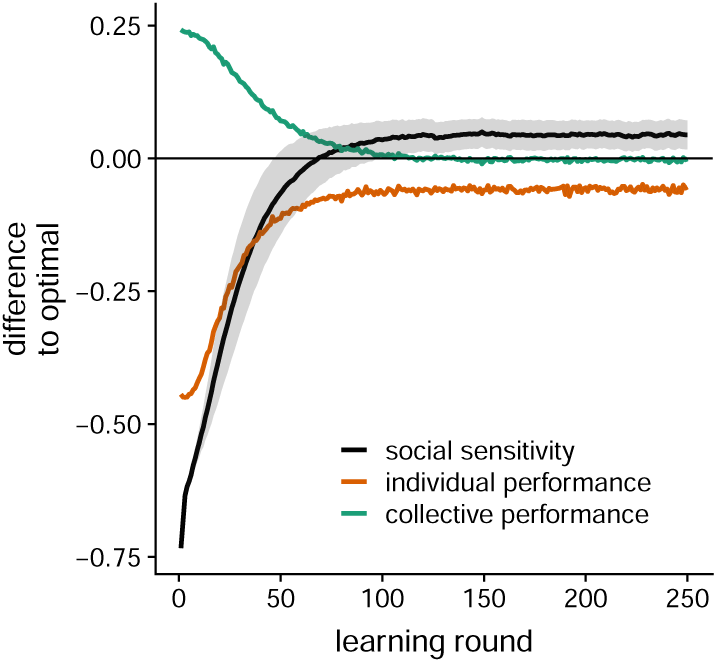
Simulation of a model in which the agents gradually update their propensity for social learning to maximize their own individual payoff (see methods for more details). The figure shows the development of the group’s average social learning level (black line, gray band shows variance) and the individual and collective performances over 150 learning rounds. Values are shown relative to a group of agents with optimal social learning *S*_*opt*_ = 0.73 (negative is lower and positive is higher than *S*_*opt*_). The group converges to a learning level slightly higher than the optimal value, causing a decay in individual performances.

## 4 Discussion

In our study, we used generic model of collective search (Gigerenzer et al., 1999; Barkoczi and Galesic, 2016; Yahosseini and Moussaïd, 2019) to address two questions. First, how does the amount of social learning impact the individual and collective performances? We find that social learning introduces a collective search dilemma. When individuals rely on social learning, their individual performances increase, whereas the collective performances decrease. We find that this dilemma can be explained by a reduction of the overall exploration volume and an increase in the ratio of consensual solutions. Social learning, therefore, is beneficial to the individual but can cause maladaptive premature convergence when individuals excessively rely on it.

Second, how does the environment moderate what is best for the individual and what is best for the group? By varying some features of the search environments, we find that social learning is a more profitable to individuals in difficult environments than in easy ones. In addition, we show that social learning is less detrimental to collective performances when all peaks are clustered in the same area.

We also studied the social environment, that is, how the features of one’s peers impact an individual’s performance. We show that any single individual can profit from a high level of social learning, but only when others are not doing so. Simulations show that this dynamics can generate a group in which individuals trying to maximize their personal payoffs by copying others end up breaking down their collective potential.

Overall, our study shows that, depending on the chosen measure of performance, the impact of social learning can change dramatically. Social learning has different effects, depending on whether one is interested in the collective performance (e.g., the group comes up with a unique collective solution (Kempe and Mesoudi, 2014) or develops a new cultural innovation (Derex and Boyd, 2016; Derex and Boyd, 2015)), or in the average individual performance (see, e.g., (Lazer and Friedman, 2007; Mason and Watts, 2012; Barkoczi and Galesic, 2016)). Here, it is interesting to point out that the measures of performance are not limited to the two instances that we have studied in this work. Other research has, for instance, considered the group’s ability to find the problem’s global optimal solution (Cooper et al., 2010; Derex and Boyd, 2016). In this case, any solution that is not the optimal one is not taken into account in the calculation of success, which comes down to considering landscapes with very low dispersion. The results of our study suggests that social learning would then have little negative effect on collective performance in that case.

Our results outline the importance of correctly balancing independent exploration and social learning (Bernstein et al., 2018; Rogers, 1988). This fact connects to a current debate about whether sparse networks that spread information slowly between individuals yield better performances than dense networks that support a rapid dissemination of information within the group.

We argue that the role of the network structure can only be understood when put in perspective with the individuals’ behavior (Barkoczi and Galesic, 2016). Sparse networks, for instance, can be beneficial when individuals tend to rely too much on each other, by regulating the flow of social information within the group (as in, Lazer and Friedman, 2007). On the contrary, dense networks are favorable when individuals have a propensity to explore independently, by supporting the rapid dissemination of a few good solutions (as in, Mason and Watts, 2012; Derex, Perreault, et al., 2018). In other words, the network structure and the behavior of the individuals are two forces that can modulate the flow of social information. A good match between them supports an efficient social dynamics during collective problem-solving.

More generally, our work helps understanding the complex dynamics that operate within interacting groups of problem-solvers and contributes to current research on collective intelligence. Future work will expand our findings to more complex problems such as NK-landscapes (Kauffman and Levin, 1987), to concrete tasks such as the traveling salesman problem (Yi et al., 2012), or to other fields that require a tradeoff between individual and collective search (March, 1991).

## 5 Methods

We model problem-solving as a search in a landscape – a conceptual representation of a solution space (Wu et al., 2018; Yahosseini and Moussaïd, 2019). In such a landscape, each field represents one solution and is associated to one payoff. We develop a model describing how individuals search in such a landscape when trying to maximize their own payoff while at the same time observing other people’s solutions. This model extends previously validated approaches for independent (Yahosseini and Moussaïd, 2019; Gigerenzer et al., 1999) and social search (Laland, 2004).

### 5.1 Search model

The individual search behavior is composed of (1) a social search rule, and an independent search rule. We assume that the agent relies on the independent search rule during a certain number of simulation rounds at the beginning of the search process, and then switches to the social search rule. The fraction of rounds during which the social search rule is applied is given by the social learning parameter *S*. For example, a value of *S* = 0.8 indicates that the agent applies the independent search rule during the first 20% of the rounds, and applies the social search rule for the remaining 80%.

The independent and the social search rule are defined as follows:

1. Independent search rule: In every round, the agent maximizes its immediate payoff by moving to the neighboring solution that offers the highest payoff. If more than one solution fit this criteria, the agent maximizes the number of explored solutions by moving away from visited areas of the landscape. If multiple solutions are still equivalent, the agent chooses among them at random. If all neighboring solutions offer a lower payoff than the current solution (i.e. if the agent is occupying a local optimum), then the search stops.
2. Social search rule: In every round, the agent first looks at the current solutions of all its peers. The solutions that cannot be reached within the remaining time are ignored. If one of these solutions offers a higher payoff than the agent’s current solution, it moves one step towards that solution, following the shortest path. If multiple solutions are on the shortest path the individual rule is applied to those. If none of one’s peers’ solutions meet these criteria, the agent resorts to the independent search rule.

### 5.2 Search environment

We implement the search environments as a landscape with multiple local optima (also denoted as “peak”). We use the following procedure to generate these landscapes (Yahosseini and Moussaïd, 2019; Wu et al., 2018):

1. We first generate 32 sub-landscapes. Each sub-landscape consists of a 99 × 99 matrix filled with zeros and will serve as layers for each of the 32 peaks. We select one random coordinate *P* around which all the peaks will be clustered.
2. For each of the 32 sub-landscapes, we draw one coordinate from a normal distribution with *mean* = *P* and *SD* = *d* that will serve as a location for the peak *p*. The parameter *d* determines how many of the 32 peaks are clustered around *P*.
3. The height of the peak *p* (i.e., corresponding to the payoff of that solution) is drawn from a normal distribution with *mean* = 0 and *SD* = 1, and is subsequently squared to avoid negative payoffs.
4. To create a local gradient around the peak *p*, we apply a Gaussian filter with *SD* = *f* (*w*) on each sub-landscape. Here *w* controls how much the width of the peak *p* and the payoff correlate. The function *f* (*w*) is defined as *f* (*w*) = (1 -*w*) *\* random* + *w × payoff* (*p*), where *random* is a random number between 1 and *payoff* (*p*). That is the width of a peak *p* is determined by a combination of its payoff and a random factor controlled by *w. f* (*w*) is linearly scaled between 1 and 4 over all peaks in the 32 sub-landscapes, resulting in a width of the peaks between 1 and 4.
5. The 32 sub-landscapes are merged into a single one by selecting the highest payoff across all sub-landscapes at each coordinate.
6. Finally, all payoffs are linearly scaled between 0 and 100.

This procedure generates landscapes similar to those shown in figure 2A.

### 5.3 Simulation procedure

For each simulation run, we randomly position ten agents in the landscape within an area of size 10 × 10 solutions. Each simulation run lasts 30 rounds. In each round, the agents behave according to the previously described model. The size of the landscape, the starting solution and the total number of rounds are balanced in such a way that the “borders” of the landscape can never be reached reached. At the end of each run, the payoff of each individual agent is the payoff associated to the current occupied solution. We report the individual performance as the groups mean payoff and the collective performance as the highest payoff found by any member of the group. For better comparison, we generally report performance measures relative to the performance of a group of non-interacting agents, that is, a group of agents with no social learning (*S* = 0). A performance value higher than one indicates a better performance compared to groups of non-interacting agents.

### 5.4 Simulating repetitive updates in social learning

In the simulations presented in figure 5, we describe how a specific social learning level could develop. For that, we assign an individual social learning level *S*_*i*_ and a direction parameter *D*_*i*_ to each agent. The direction parameter *D*_*i*_ describes the direction in which the social learning level of that agent is changed after each learning round (i.e., whether it increases or decreases). In each learning round, all agents search 50 landscapes as described in the simulation procedure. At the end of this process, each agent compares its average payoff to the payoff obtained in the previous learning round. If the payoff has decreased, the agent switches the direction *D*_*i*_ (i.e., from increase to decrease and vice versa). Afterwards, all agents change their social learning level by one step according to *D*_*i*_ and a new learning round starts. Here, we assume that all agents start from a non-social state (*S* = 0) and initially increase their social learning level. The results are robust to different parameter values.

